# Lifelong molecular consequences of high Glucocorticoids exposure during development

**DOI:** 10.1101/2023.02.13.528363

**Authors:** Min-Kyeung Choi, Alex Cook, Kanak Mungikar, Helen Eachus, Anna Tochwin, Matthias Linke, Susanne Gerber, Soojin Ryu

## Abstract

Early life stress (ELS) is one of the strongest risk factors for developing psychiatric disorders in humans. As conserved key stress hormones of vertebrates, glucocorticoids (GCs) are thought to play an important role in mediating the effects of ELS exposure in shaping adult phenotypes. In this process, early exposure to high level of GCs may induce molecular changes that alter developmental trajectory of an animal and primes differential adult responses. However, comprehensive characterization of identities of molecules that are targeted by developmental GC exposure is currently lacking. In our study, we describe lifelong molecular consequences of high level of developmental GC exposure using an optogenetic zebrafish model. First, we developed a new double-hit stress model using zebrafish by combining exposure to a high endogenous GC level during development and acute adulthood stress exposure. Our results establish that similar to ELS-exposed humans and rodents, developmental GC exposed zebrafish model shows altered behavior and stress hypersensitivity in adulthood. Second, we generated time-series gene expression profiles of the brains in larvae, in adult, and upon stress exposure to identify molecular alterations induced by high developmental GC exposure at different developmental stages. Third, we identify a set of GC-primed genes that show altered expression upon acute stress exposure only in animals exposed to a high developmental GC. Interestingly, our datasets of GC primed genes are enriched in risk factors identified for human psychiatric disorders. Lastly, we identify potential epigenetic regulatory elements and associated post-transcriptional modifications following high developmental GC exposure. Thus, we present a translationally relevant zebrafish model for studying stress hypersensitivity and alteration of behavior induced by exposure to elevated GC levels during development. Our study provides comprehensive datasets delineating potential molecular targets underlying the impact of developmental high GC exposure on adult responses.

## Introduction

Early life stress (ELS) affects healthy development and aging as well as susceptibility to psychiatric disorders in humans^1–4^. Studies in humans and animals suggest ELS may increase risk for psychiatric diseases such as depression by inducing hypersensitivity to future stress exposure^5–8^. However, mechanism by which ELS sensitizes responses to future stress responses in adulthood remains poorly understood. In vertebrates, stress response is mediated by a highly conserved neuroendocrine system called the Hypothalamic-Pituitary-Adrenal axis (HPA-axis) in humans^9^ and the Hypothalamic-Pituitary-Interrenal axis (HPI-axis) in fish whose activation result in the release of Glucocorticoids (GCs)^10^. While GCs have pleiotropic effects on many aspects of animal physiology, exposure to elevated GCs during brain development is known to cause long-term alteration of behavior, physiology, and stress regulation^11^. However, how exposure to high levels of GC during development predisposes individuals to adulthood dysfunction is poorly understood. One potential mechanism is long-lasting epigenetic changes, which modify how animals respond to stress in adulthood^12^. For example, altered DNA methylation at the glucocorticoid receptor (GR) gene promoter induced by poor maternal care is associated with adulthood alterations in histone acetylation, DNA methylation, transcription factor binding, GR expression, and the HPA response to stress^13^. Moreover, a recent study identified differentially expressed genes (DEGs) “primed” by ELS in specific regions of ELS-exposed mouse brains^14^. These genes showed enhanced differential expression in ELS-exposed animals in adulthood upon re-exposure to stress and may be associated with behavioral changes in adulthood. Another study identified direct GC-primed DEGs and associated long-lasting DNA methylation alterations using a GC (dexamethasone)-exposed human hippocampal progenitor cell line^15^. Together, the above studies establish that ELS or developmental GC exposure results in distinct responses to stress later in life by priming and maintaining longer-term alteration of specific genes. However, so far, ELS- or GC-primed genes have only been identified in specific regions of the brain including reward circuitry or specific cell lines, and brain-wide GC-primed molecular alterations have not been investigated.

Here we report whole brain transcriptomic alterations of high GC-exposed fish across the life course and upon acute stress exposure in adulthood. Interestingly, when subjected to acute stress in adulthood, developmental high GC-exposed fish show highly exaggerated endocrine and transcriptional responses, indicating that adult stress is processed differently depending on the GC exposure history of an animal. The genes primed by developmental GC-exposure were enriched in gene sets associated with human neuropsychiatric disorders, suggesting similarity of molecular mechanisms by which early GC exposure leads to altered adult functions among vertebrates. We identify hitherto uncharacterized novel GC-primed genes that are enriched in synapse and neuronal signaling function. Whole brain DNA methylation analysis identified some of them as direct targets of developmental GC-mediated epigenetic modifications. Lastly, the expression of many epigenetic modulators affecting RNA processing, histone modification and DNA modification are altered following acute stress in developmental GC-exposed individuals, which can contribute to the different physiological and behavioral responses following adult stress exposure in later stage of life.

## Results

### Double-hit zebrafish stress model exhibit altered adulthood stress response

We developed a double-hit stress model using zebrafish which combines exposure to a high level of GC during development with acute adult stress exposure (Fig. 1a). Firstly, we used our previously reported optogenetic transgenic model *Tg(star:bPAC-2A-tdTomato)* where the elevation of endogenous cortisol (GC in fish) is induced by blue light which activates *beggiatoa* Photoactivated Adenylyl Cyclase (bPAC), expressed specifically in steroidogenic interrenal cells^16–18^ (Fig. 1b). Like the mammalian adrenal gland, the fish interrenal gland is composed of two cell types: aminergic chromaffin cells and steroidogenic interrenal cells, the latter of which produce glucocorticoids (GCs). We established that *Tg(star:bPAC-2A-tdTomato)^uex300^* raised under ambient light containing blue light (hitherto referred to as bPAC+) showed increased cortisol level and expression of *fkbp5*, one of known GR signaling marker genes^19,20^, at larval stages^17,18^ (Fig. 1c). Elevated cortisol levels through larval and juvenile stages were detected in bPAC+, but not maintained until adulthood (after 3 months) allowing us to achieve a robust and persistent increase in endogenous cortisol levels during development (Fig. 1d-e). Throughout the study, bPAC+ fish were compared to bPAC-, siblings of bPAC+ which do not carry the transgene themselves but are offspring of a bPAC+ parent crossed to wild type. We also compare their phenotypes to wildtype TU, which is the same strain as bPAC+ and bPAC-, to determine possible effects resulting from elevated GC in parental generation (S. Fig. 1). Secondly, for acute stress delivery, we utilized a looming dot stimulus (LD), which mimics an approaching predator^21^ (Fig. 1f). bPAC+ adults exhibited an exaggerated endocrine response after LD exposure compared to wildtype or bPAC-adults and even an elevated GC level induced by handling before the LD exposure (Fig. 1g), indicative of a highly sensitized endocrine response to acute stress in bPAC+.

**Figure 1.**
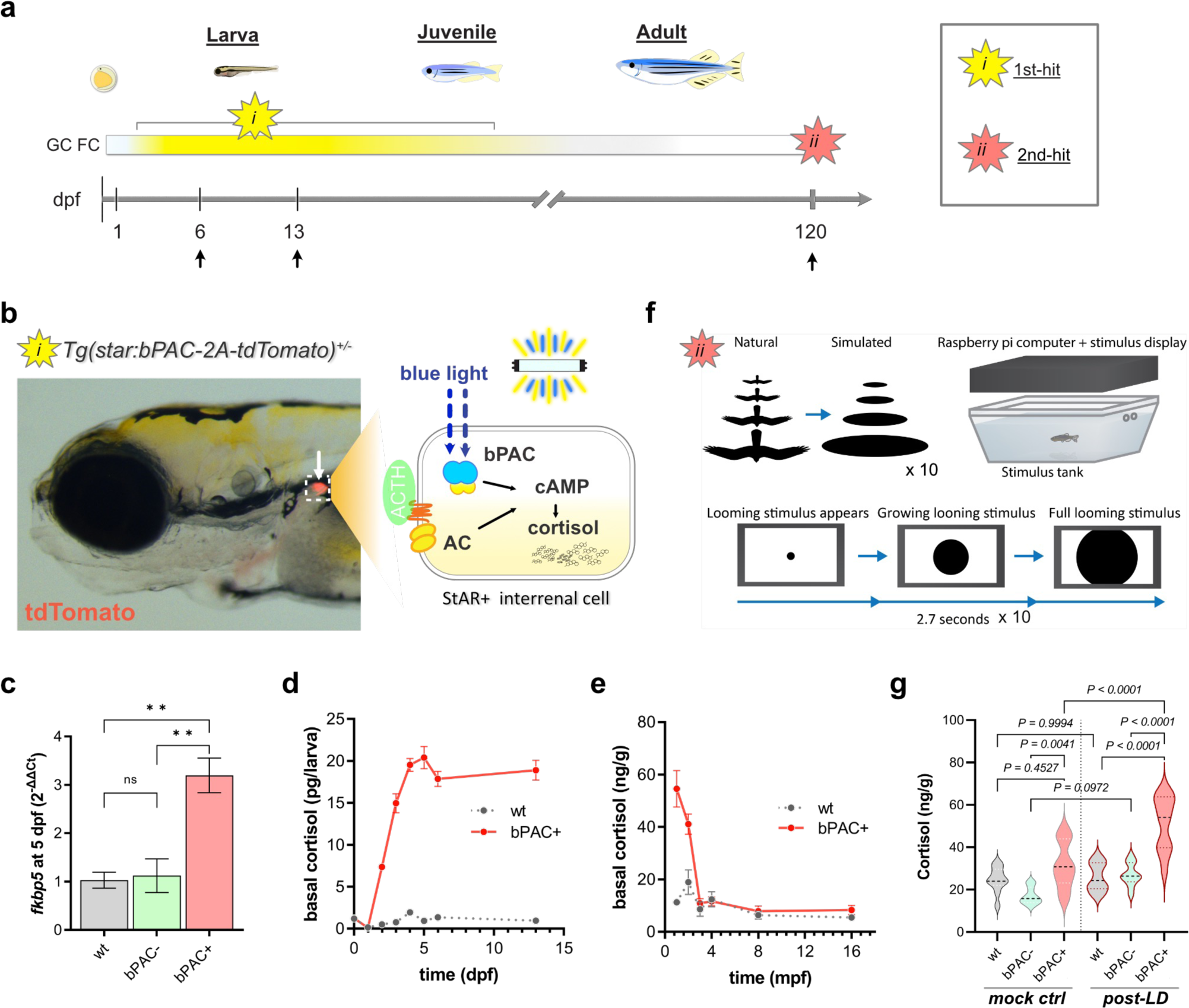
Double-hit stress model shows elevation of cortisol during development and upon acute stress exposure in adulthood. (a) A scheme illustrating the strategy for underlying Double-hit zebrafish stress model which combines *i)* optogenetic cortisol elevation during development and *ii)* adulthood acute stress exposure. Arrows indicate sample collection times points. (b) An image of *Tg(star:bPAC-2A-tdTomato)^+/-^*larva at 5 dpf. The transgene is expressed solely in the interrenal gland (arrow). A schematic drawing representing the intracellular events within the steroidogenic interrenal cells of the transgenic fish (dotted box). Endogenous GC production is stimulated by ACTH signaling. bPAC mimics ACTH signaling by increasing cAMP when exposed to blue light. (c) One of the well-known GR signaling markers, *fkbp5*, was significantly upregulated in bPAC+ larvae compared to bPAC- and wildtype control. ** = *P*<0.01. Tukey’s multiple comparisons test following one-way ANOVA, F (2, 6) = 16.25, *P*=0.0038. (d-e) Basal cortisol level at larval stages and from juvenile to adult. (f) Schematic of the acute stress paradigm. Looming dots mimic approaching predators from above (Cook *et al*. 2023). Total 10 times of LD were given as an acute stress. (g) bPAC+ showed significantly higher cortisol levels following LD, compared to handling controls (no LD presented on the screen). Tukey’s multiple comparisons following one-way ANOVA; F(5, 54)=19.31, *P*<0.0001. Abbreviations: dpf: day post fertilization, mpf: month post fertilization, GC: glucocorticoid, ACTH: Adrenocorticotropic Hormone, AC: adenylyl cyclase, cAMP: Cyclic adenosine monophosphate, FC: fold change, LD: Looming dots, wt: wild type, ctrl: control, bPAC: *beggiatoa* photoactivated adenylyl cyclase.

### High GC-exposed animals during development show altered behaviors in adulthood

To determine whether exposure to a high level of endogenous GC during development (dGC) alters adult functions, we tested adult behavior encompassing different behavioral domains using wildtype, bPAC- and bPAC+ (Fig. 2). We first established that they did not show significant differences in basal locomotor activity (Fig. 2b-c). We then used a novel tank test to assess their adaptive responses in a novel environment^22,23^ (Fig. 2d-g). When placed in a novel tank, zebrafish typically first dive to the bottom whilst performing bouts of erratic high-speed swimming^22,23^. There was no significant difference between bPAC- and wildtypes in average depth or fast swimming time (Fig. 2e, g). In contrast, bPAC+ fish showed reduced depth-preference (Fig. 2e) and reduced fast-swimming compared to bPAC- and wildtype fish (Fig. 2g). In addition, bPAC+ showed reduced average speed compared to wild type (Fig. 2f). Next, we tested feeding behavior by counting the number of floating food pellets eaten by a fish during a 10-minute interval (Fig. 2h). Wild-type and bPAC-adult fish consumed on average 14.5 and 11.25 pellets, respectively while bPAC+ fish consumed significantly fewer pellets (on average 2.33 pellets out of 25) (Fig. 2i-j). This was not due to the lack of recognition nor interest in the food, as bPAC+ fish approached the food pellets and spent a longer time near the pellets (feeding zone) than bPAC- and wildtype fish (Fig. 2k). Further, we analyzed social behavior using a new test that we recently established, which allows measuring both social approach and social interaction within a same paradigm (Fig. 2l). While bPAC+ exhibited reduced social interactions compared to wild type, no significant difference was observed in bPAC+ compared to bPAC- (Fig. 2m-o). Similarly, whilst bPAC+ exhibited altered fear conditioning behavior compared to wildtype, there was no significant difference observed between bPAC+ and bPAC- in an associative learning test using a Pavlovian avoidance learning test^24,25^ (Fig. 2p-s). Thus, exposure to a high level of endogenous GC during development led to an exaggerated endocrine response upon acute stress exposure in adulthood and alteration in responses in novel environment and feeding behaviors. For social and fear conditioning behaviors, exposure to a high GC during parental generation appear to be sufficient to modify adult behavior.

**Figure 2.**
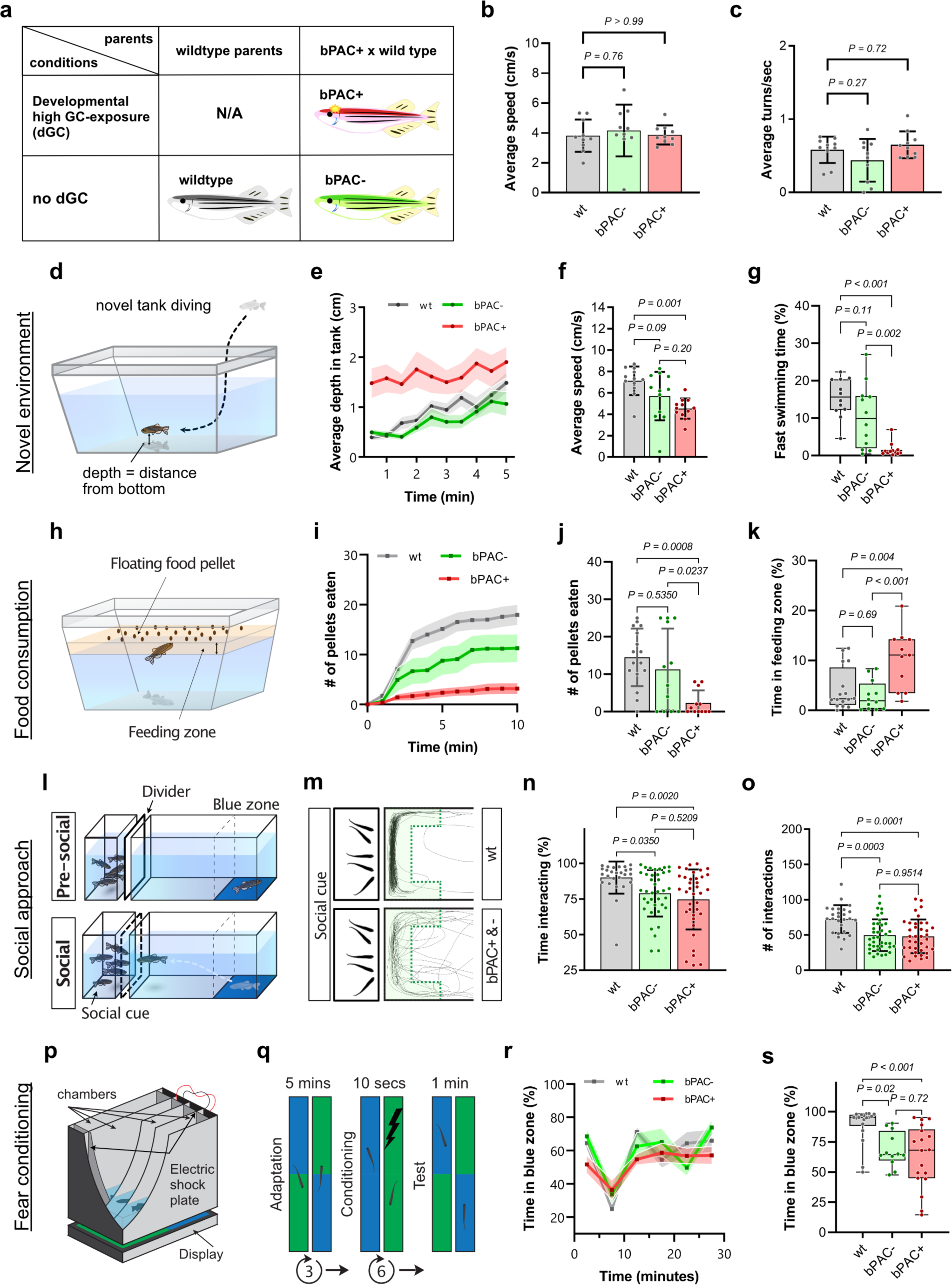
Altered behaviors in adult bPAC+, bPAC- and wildtype fish. (a) A schematic depicting experimental groups. (b-c) No significant differences were observed in average speed nor turn frequency among wild type, bPAC+ and bPAC-. Dunnett’s multiple comparisons followed one-way ANOVA, b) F(2, 27)=0.2248, *P=0.80*; c) F(2, 27)=2.332, *P=0.12*. (d) A schematic drawing for the novel tank test. (e) Average depth as measured by average distance from the bottom of tank was greater for wild type and bPAC-compared to bPAC+. (f-g) bPAC+ fish showed a slower average speed than wild type and reduced fast swimming bouts compared to bPAC- and wild type. (h) A schematic for the feeding behavior test. (i-j) bPAC+ significantly consumed fewer number of pellets within 10 min compared to wild type or bPAC-. (k) bPAC+ spent more time near the food compared to wildtype or bPAC-. (l) Schematic figures for the social approach and maintenance test. (m) Representative traces of wild-type, bPAC+ and bPAC-when conspecifics are visible. The green shaded area indicates the interaction zone where wildtype fish typically spent most of their time when conspecifics are visible. (n-o) During this social phase, bPAC+ and bPAC-showed a significantly reduced duration and frequency of social interactions compared to the wild type (Cook rt al., 2023}. (p) A schematic drawing for a chamber used for the fear conditioning test. (q) An electric shock was given to subject fish when they swim in the green colored area. After conditioning, time spent in each color zone was measured. (r) No significant difference was observed in color preference prior to fear conditioning. (s) bPAC+ showed less preference for the safe zone color (blue) after the conditioning using an electric shock. Error bars on bar graphs indicate the mean±SD and shading in line graphs represents the mean±SEM. Tukey’s multiple comparisons followed one-way ANOVA. f) F(2, 33)=7.806, *P=0.002*; g) F(2, 33)=17.31, *P<0.0001*; j) F(2, 37)=8.364, *P=0.001*; k) F(2, 36) = 9.120, *P<0.001*, n) F(2, 98)=6.273, *P=0.0027*; o) F(2, 98)=10.93, *P<0.0001*; s) F(2, 45)=8.927, *P<0.001*.

### Distinct transcriptional alterations in bPAC+ fish across the life course

To identify transcriptional changes caused by dGC-exposure, we performed time series whole-brain RNA sequencing using larval (6 and 13 dpf) and female adult (120 dpf) brains of bPAC+, bPAC-, and wildtype fish. The total number of DEGs identified among different genotypes varied starkly at different time points. The global gene expression profiles of bPAC+ were clearly distinct from those of wild types while bPAC-profiles mapped more closely to those of bPAC+ than wild type (S. Fig. 2a). The overall expression pattern of identified DEGs using the threshold (FDR < 0.05 and |log_2_(fold-change, FC)| > 1.5, S. Data File 1) mainly exhibited a close grouping primarily based on groups and time point except for two 13 dpf bPAC-profiles (Fig. 2a).

In bPAC+ and bPAC-comparisons which reveal alterations induced by dGC-exposure, a large number of DEGs were identified at the early larval stage (6 dpf, up/downregulated DEGs: 1147/1631), but a much fewer number of DEGs were detected at 13 dpf (28/79) and 120 dpf (47/278). At 6 dpf, we observed transcriptional alterations of genes that are previously reported to be altered following GC treatment, including *fkbp5*^19,20^ and *hsd11b2*^26^ (Fig. 3b). In the adult bPAC+ brains, 27% (13 genes) of the upregulated and 31% (88 genes) of the downregulated DEGs overlapped with DEGs from larval stages (Fig. 3c).

**Figure 3.**
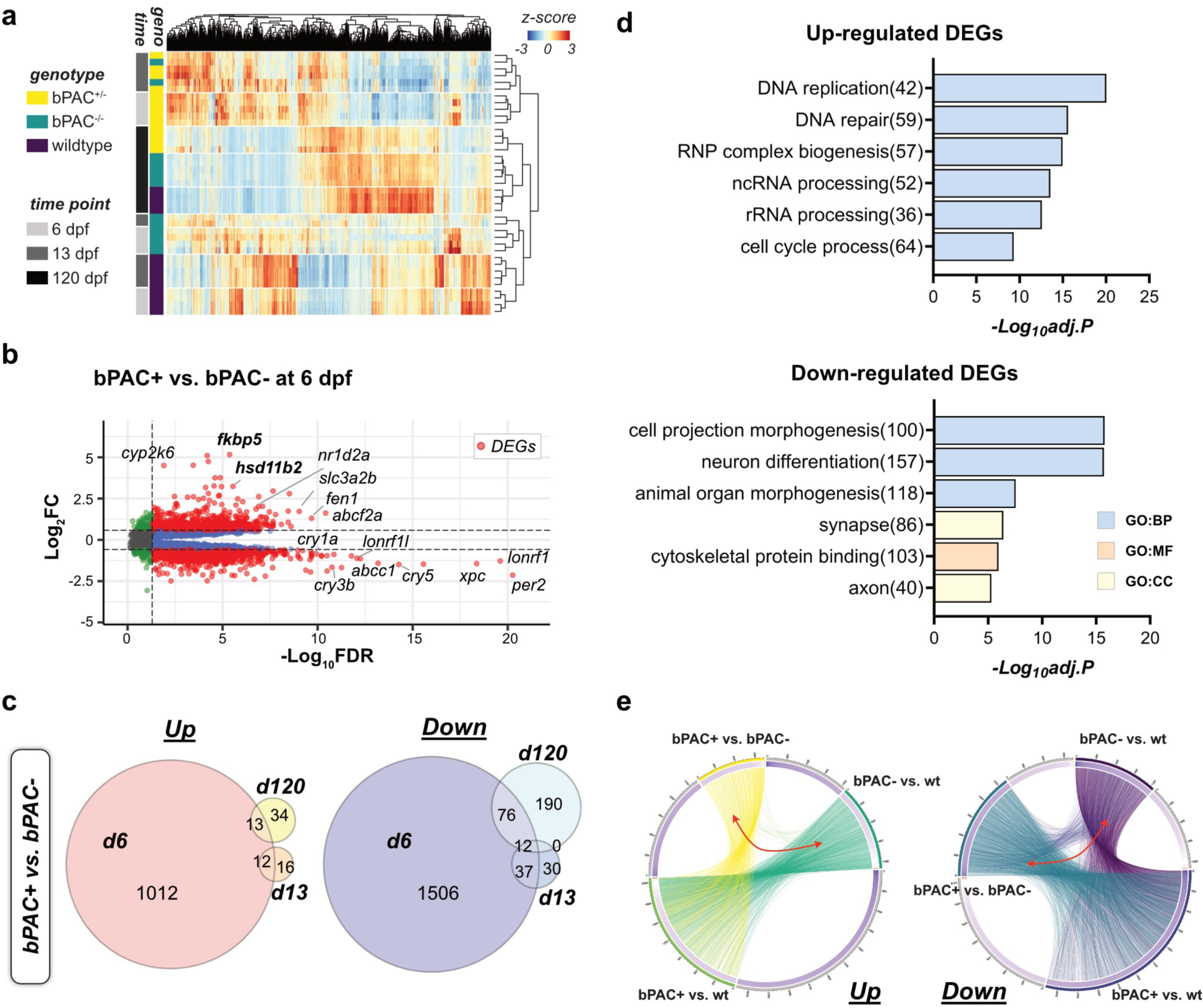
Developmental GC exposure leads to differential gene expressions across the life course. (a) Read counts of genes by genotype and time points are represented by z-score in the heat map. Bars on the left side indicate the genotype of fish (yellow: bPAC+, green: bPAC-, dark purple: wild type) and time points (gray: 6 dpf, dark gray: 13 dpf, black: 120 dpf). (b) Volcano plot showing DEGs (red dots) in bPAC+ and bPAC-brain transcriptome at 6 dpf. Dashed lines indicate FDR and FC cut-off values set at 0.05 and 1.5, respectively. Labels identify the genes associated with cortisol or circadian rhythm among the smallest FDR values. Plots for other time points are available in the Supplementary data file. Complete list of identified DEGs described in S. Table 1. (c) Venn Diagram for up- and down-regulated DEGs among wild type, bPAC- and bPAC+; criteria for DEG identification: |FC|>1.5 and FDR <0.05. (d) Top enriched GO terms for up- and down-regulated DEGs in bPAC+ compared to bPAC- at 6 dpf. Complete list of DEG-enriched GO terms are described in S. Table 2. (e) Circos plots showing overlapping DEGs by linked line among three pairwise comparisons, bPAC+ vs. bPAC-, bPAC+ vs. wild type, bPAC- vs. wild type. FDR: False discovery rate, GO: Gene ontology, *adj.p*: adjusted *P*-value.

Next, we performed GO enrichment tests comparing bPAC+ and bPAC-. Upregulated DEGs in bPAC+ fish brains at 6 dpf were enriched in RNA processing and DNA replication and repair (upper panel of Fig. 3d, S. Table 2). In contrast, downregulated DEGs at 6 dpf were enriched in processes related to nervous system development including neuron differentiation and cell morphogenesis (bottom panel of Fig. 3d, S. Table 2). Indeed, we confirmed alteration of neurogenesis in larval bPAC+ fish^18^. Together these results indicate dysregulation of normal developmental processes in bPAC+ brains in early life coupled with potential genomic instability, replication stress^27^, and delayed or immature differentiation of neuronal cells. At 120 dpf, despite a smaller numbers of DEGs identified (Fig. 3c), we noted significant upregulation of genes involved in pre-mRNA splicing and 5-hydroxymethyl cytosine (hmC) reading (*wdr76*, *neil1*, and *hells*)^28^ suggesting differential epigenetic processes while downregulation of some of the marker genes for oligodendrocyte (*sox10*, *olig2*) indicating decreased oligodendrocyte differentiation in adult bPAC+ brains (S. table 1).

Further, we carried out bPAC+ and wild type or bPAC- and wild type comparisons to determine potential effects resulting from high GC exposure that occurred in the bPAC+ parental generation. Even with more stringent thresholds to identify DEGs (FDR < 0.01 and |log2FC| > 2) than those of previous comparisons, the upregulated DEGs enriched GO terms in later developmental stages were similar to those identified in the bPAC+ vs. bPAC-comparison including cell cycle, DNA metabolic process, and RNA processing processes (left panel of S. Fig. 2c). Downregulated DEGs across all time points were enriched in processes related to cell adhesion and cell migration, as well as nervous system development including neuron differentiation, cell morphogenesis, and axon development (right panel of S. Fig. 2c), similar to the bPAC+ versus bPAC-comparison. Also, the identified upregulated DEGs in the larval stage of bPAC+ compared to wild type were enriched in potential metabolic enzymes for GC such as cytochrome monooxygenases, 11β-hydroxysteroid dehydrogenase or sulfotransferases^29–31^ (left panel of S. Fig. 2c). Some of those DEGs were also significant even in the comparison between bPAC- and wildtype (S. Table 2) suggesting alteration in GC metabolism resulting from elevated GC in parental generation. Lastly, 2.08% (27/1295) and 16.6% (211/1259) of up- and downregulated DEGs identified during larval stages in the comparison between bPAC- and wild type overlapped with DEGs identified in the comparison between bPAC+ and bPAC- (Fig. 3e, marked with red double-headed arrows, S. Fig. 2d). However, in adulthood, much fewer overlapping DEGs were identified compared to larval stages (S. Fig. 2d). Together these results identify transcriptional differences caused by dGC exposure and suggest the existence of alterations induced by high GC exposure in the parental generation apparent in both bPAC+ and bPAC-.

### High GC exposure during development leads to exaggerated transcriptional response to acute stress in adulthood

We next examined transcriptional response of bPAC+ upon acute stress exposure in adulthood. To identify a class of genes that are induced by acute stress exposure only in bPAC+, we carried out comparisons by genotype (bPAC+ vs. bPAC-) or condition (pre- vs. post-LD) (Fig. 4a). bPAC+ fish showed a much greater transcriptional response following LD exposure compared to bPAC-fish (Fig. 4b). Strikingly, bPAC+ showed more than 17-fold greater number of DEGs in response to LD exposure (bPAC+ in Fig. 4c, 1754 genes) compared to bPAC- (bPAC- in Fig. 4c, 102 genes). 83% of DEGs induced by LD-exposure in bPAC- were identical to those in bPAC+, while only 17 DEGs were specific to bPAC-. These results indicate that dGC- exposure primed a large number of latent alterations which become only apparent upon acute stress exposure in adulthood in bPAC+ fish. We refer to these differentially regulated genes following LD-exposure in bPAC+ (bPAC+ or bPAC- specific DEGs; 1669 and 17 genes) as developmental GC-primed genes (dGC-primed genes, a total of 1686 genes) (Fig. 4c). dGC- primed genes were significantly overrepresented in the GO terms synapse, cell morphogenesis, regulation of signaling, and more (Fig. 4d-e, S. Table 4), and approximately 80% of dGC- primed genes (1362 genes) were downregulated. For example, synapse associated dGC-primed genes were mostly downregulated and showed enhanced alteration following LD (Fig. 4e).

**Figure 4.**
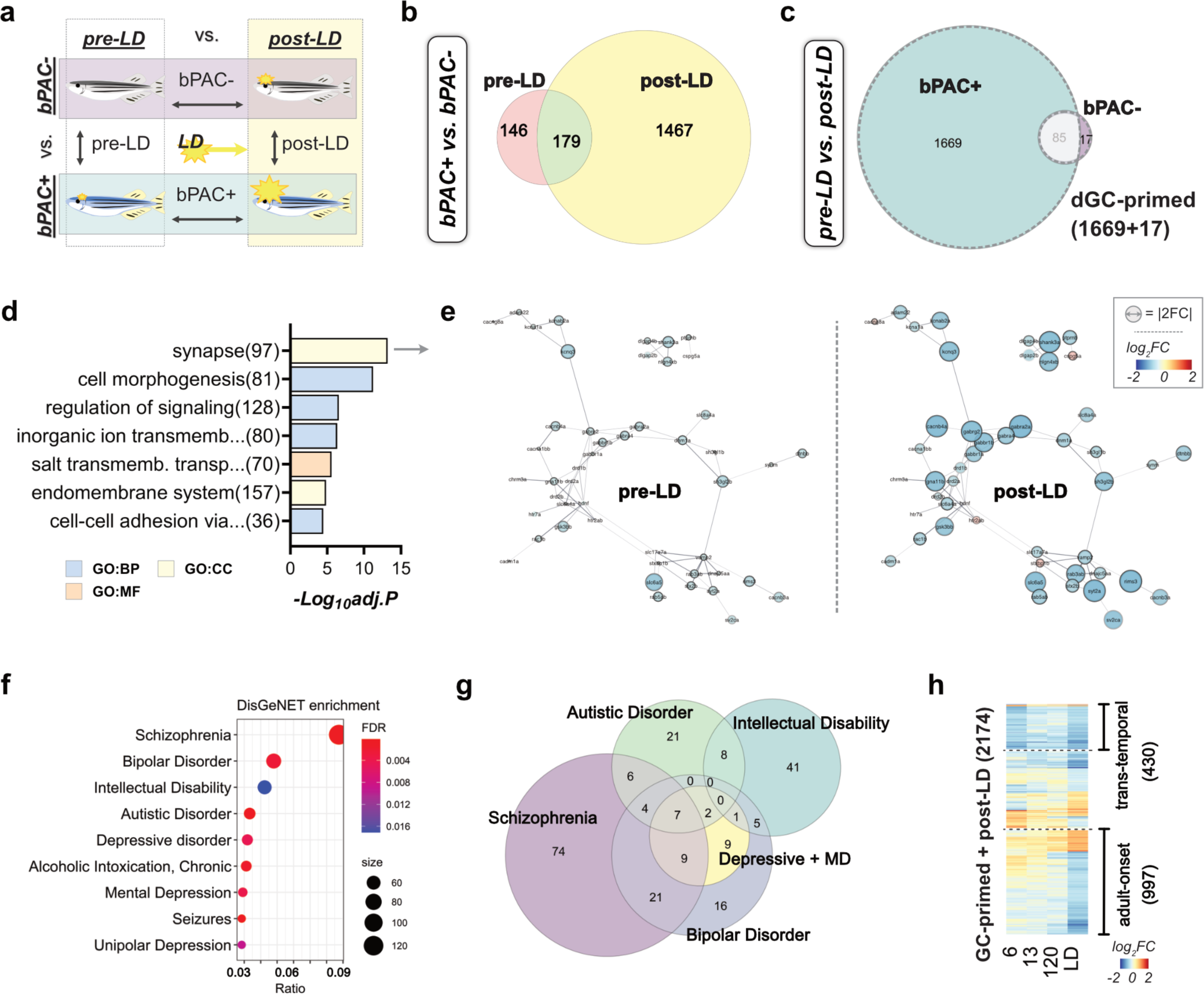
Developmental GC exposure primes differential gene expression following acute stress in bPAC+ adults. (a) Venn diagram representing DEGs between bPAC+ and bPAC- in pre-LD and post-LD conditions. (b) Schematic for three comparisons to identify transcriptional changes in response to LD exposure in adults. DEGs were determined by FC and FDR. |FC|>1.5 and FDR<0.05. (c) Venn diagram showing post-LD DEGs from bPAC+ and bPAC-. The list of DEGs is described in S. Table 3. dGC-primed genes are marked by dashed line. (d) The top GO terms enriched in dGC-primed genes. The enrichment score = −log_10_(adjusted *P*-value). (e) PPI networks of synapse associated dGC-primed genes. The size of node indicates fold-change. Networks sourced by STRING. (f) Result of human disorders enrichment test based on the DisGeNET database using GC-primed genes. The ratio represents overlapping/input genes. Top ranked disorders are Schizophrenia (FDR= 1.12e-04, ratio=122/883), Bipolar Disorder (FDR= 2.47e-03, ratio=69/477), Intellectual Disability (FDR= 1.71e-02, ratio=61/447), Autistic Disorder (FDR= 1.87e-04, ratio=48/261), Depressive disorder (FDR= 4.51e-03, ratio=46/289), and Mental Depression (FDR= 4.06e-03, ratio=42/254). The complete results are noted in S. Table 5. (g) Venn diagram for dGC-primed genes that are associated with the top human psychiatric disorders in panel f. (h) Heatmap showing log2FC of 2174 LD-DEGs across time points demonstrated 430 trans-temporal and 997 adult-onset genes were mostly downregulated. LD: looming dots, FC: fold change, FDR: False discovery rate, PPI: Protein-protein interaction, ratio: GC-primed genes/disease genes, Depressive + MD: Depressive disorder and Mental Depression.

Among the genes associated with GC signaling, we observed that the expression of *nr3c1* (GR) is not altered in bPAC+ adults while the expressions of *nr3c2* (mineralocorticoid receptor; MR) and *crhr1* (corticotropin releasing hormone receptor) were significantly downregulated only in bPAC+ adults following LD exposure (S. Fig. 3a). Moreover, we identified that several key molecules within the CRH signaling pathway were altered in adult bPAC+ brains following LD (S. Fig. 3b), consistent with the observed downregulation of *crhr1* in bPAC+ following LD (S. Fig. 3a).

### dGC-primed genes are associated with neuropsychiatric disorders in humans

Since ELS is a strong risk factor for developing psychiatric disorders in humans which manifest in adulthood triggered by stressful life events^2–4^, we asked whether dGC-primed genes could overlap with psychiatric disease risk factors identified in humans. Remarkably, human homologs of dGC-primed genes exhibit a significant enrichment of known psychiatric disease-associated genes curated in the DisGeNET database^32^ (Fig. 4f, S. Table 5). Top-ranked psychiatric disorders were schizophrenia, bipolar disorder, intellectual disability, autistic disorder, and depressive disorder (Fig. 4f-g). In addition, we carried out a comparison with a recent study that reported 702 GC-primed DEGs linked with long-lasting differentially methylated sites following re-exposure to GC (dexamethasone) in a human hippocampal progenitor/neuron cell line^15^ and identified 103 genes overlapping with DEGs identified in post-LD bPAC+ vs. post-LD bPAC- comparison (up/down=58/45) and 19 of those were dGC- primed genes (S. Fig. 4, S. Table 6). Notably, some of these post-LD bPAC+ DEGs were already modified at 6 dpf in bPAC+ (transtemporal, 430 DEGs), even though the magnitude of alterations were not always maintained at different stages while some others were altered uniquely in bPAC+ adulthood upon acute stress (adult-onset, 997 DEGs) (Fig. 4h). These results suggest that some dGC-primed genes can be already predicted in early stages following dGC-exposure. Thus, our result identifies a hitherto uncharacterized set of dGC-primed genes that are overrepresented in synapse and cell signaling molecules as well as homologs of human psychiatric disease risk factors.

### A subset of GC-primed transcriptional alterations is associated with differential DNA methylation and alternative splicing patterns in adulthood

To identify potential mechanisms establishing dGC-primed transcriptional alterations, we analyzed expression patterns of DNA methylation modifiers across different time points (S. Fig. 5). Among them, we found a significant difference in the levels of *dnmt3aa* and *dnmt3bb.3* in the bPAC+ fish compared to bPAC- and wild type at 6 dpf (Fig. 5a). The significant difference in *dnmt3aa* level in bPAC+ was maintained into adulthood. Similarly, the level of *tet1,* implicated in demethylation, is altered in bPAC+ adults compared to bPAC- and wildtype after LD exposure (Fig. 5a).

**Figure 5.**
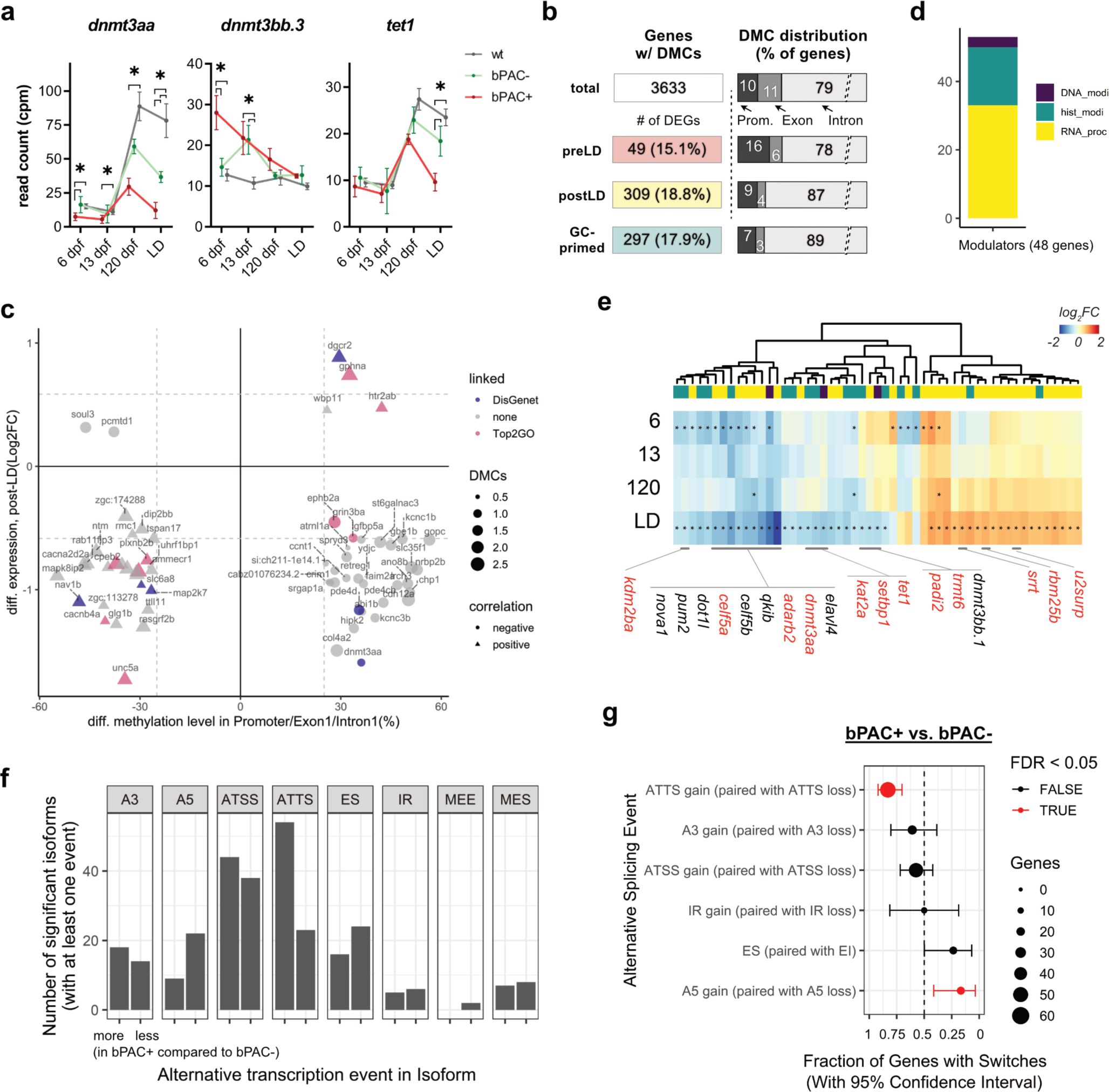
Subsets of GC-primed genes are differentially methylated and spliced in bPAC+ brains. (a) Expression trajectories for differentially expressed *dnmt*s in bPAC+ brains. Šídák’s multiple comparisons test followed mixed model ANOVA. The asterisks indicates the time points with significant differential expressions (*adj.P* <0.05 and |FC| >1.5). *dnmt3aa*: F(6, 39) = 13.13, *P<0.0001*; *dnmt3bb.3*: F(6, 39) = 11.26, *P<0.0001*; *tet1*: F(6, 39) = 7.037, *P<0.0001*. (b) The bar graph depicts the number of DEGs exhibiting statistically significant differences in DNA methylation levels, specifically on CpG sites, within both their promoter and gene body regions. (c) Methylation levels of DMCs on the regulatory regions of genes including promoter, exon1, and intron1 were positively or negatively correlated with subsets of GC-primed genes. (d) Bar graph showing known epigenetic regulators associated with DNA (purple, 3), RNA (yellow, 30), and histone (dark green, 15) modification or process among dGC-primed genes and post-LD DEGs. (e) Heatmap represents log_2_FC of them. The asterisk in each cell refers to FDR < 0.05. Red label indicates existence of DMCs on their corresponding regions. (f) Summary of identified alternative splicing events between bPAC+ vs. bPAC-. (g) Enrichment test of alternative splicing events. ES: Exon Skipping, MEE: Mutually Exclusive Exon, MES: Multiple Exon Skipping, IR: Intron Retention, A5: Alternative 5’end donor site, A3: Alternative 3’end acceptor site, ATSS: Alternative Transcription Start Sites, ATTS: Alternative Transcription Termination Sites.

To determine whether differential DNA methylation contributes to dGC-primed transcriptional alterations following LD exposure, we identified methylated cytosines (5mCs) in the adult brains of bPAC+ before LD exposure using whole genome sequencing via Oxford nanopore sequencing (S. Fig. 6). Since the majority of dGC-primed genes defined here (1657 genes) are differentially regulated in bPAC+ compared to both wild type and bPAC- (S. Fig. 6a) and previously published reports described transgenerational effects on DNA methylation in zebrafish^33,34^, we carried out a comparison of bPAC+ with wildtype brain to identify differentially methylated CpG sites (DMCs). 17.9% of dGC-primed genes (297/1657 genes) exhibited significant DMCs within their promoter and/or gene body regions (Fig. 5b). Furthermore, we identified a reduced ratio of dGC-primed genes with DMCs in their exonic regions compared to the overall distribution for all DEGs (Fig. 5b). For further analyses, we first focused on the 4937 DMCs which are located within regulatory regions of 64 GC-primed genes (promoter, exon1 and intron1)^35^ (S. Table 7). Among them, 2624 and 2313 DMCs on 33 and 31 GC-primed genes showed a negative or correlation between DNA methylation level and gene expression, respectively (Fig. 5c). Although common biological functions encompassing these 64 GC-primed genes were not found, we observed genes associated with gene expression regulators (*dnmt3aa*, *hipk2*, *ccnt1*) as well as top two GO terms in Fig. 4d (synapse and cell morphogenesis; *gphna*, *plxnb2b*, *sema5ba*, and *unc5a*) (Fig. 5c). In addition, we found hypermethylated CpGs on genomic regions associated with dGC-primed genes, *auts2a*, *mapre3a*, *igfbp5a*, *pde4cb*, which overlapped with identified GC-primed DEGs in in vitro dexamethasone treated human neurons^15^ (S. Table 7). Overall, these results suggest that altered DNA methylation patterns may associate with establishing of dGC-primed genes in bPAC+ brain.

Lastly, to identify additional mechanisms establishing dGC-primed gene expression, we analyzed the expression pattern of a broader category of epigenetic modifiers in dGC-primed and post-LD DEG gene sets. A total of 48 identified genes were associated with RNA processing including splicing and modification (yellow bar, up/down=18/12), histone modification (dark green bar, up/down=2/13), and DNA methylation modification (purple bar, up/down=0/3) which were altered following LD-exposure in bPAC+ brain (Fig. 5d). 13 genes (corresponding 27%) have at least three significant DMCs on their promoter and gene body regions suggesting a differential regulation mediated by 5mC patterns (Fig. 5e, labeled in red). Strikingly 62% of altered epigenetic modifiers we identified were associated with RNA processing including RNA splicing factors that are involved in U1, U2, or U12-dependent mRNA splicing and RNA-binding proteins that may play important roles in the nervous system^36–41^. Indeed, we identified differential usage of transcript variants between bPAC+ and bPAC- in 64 dGC-primed genes (3.8% of dGC-primed genes, Fig. 5f, S. Table 9, S. Data File 2). The most common and significant alterations in dGC-primed genes were alternative transcription termination sites (ATTSs) and alternative 5’end donor site (A5) (Fig. 5g). Together these results point to DNA methylation and RNA processing as regulatory mechanisms establishing dGC-primed transcriptional alterations.

## Discussion

We report here that a gene set enriched in the processes of DNA/RNA processing, neuron projection, and neuronal signaling components are altered following dGC-exposure. Strikingly dGC exposure led to exaggerated endocrine stress responses and primed a set of genes to exhibit altered transcriptomic responses to acute stress in adulthood. We identified a comprehensive list of brain-wide dGC-primed gene set and revealed that some of these DEGs are associated with neuropsychiatric disorders in humans and include a set of gene expression modulators. Lastly, we identify DNA methylation and RNA processing as gene regulatory mechanisms that may contribute to establishing of the dGC-primed response to acute stress in adulthood.

The dGC-primed gene set identified here shares a significant overlap with a previous report of GC-primed genes using neuronal cells derived from human^15^. However, we could only confirm a limited number of overlapping DNA methylation pattern alterations between our study and those identified in the previously published study. DNA methylation patterns and gene expression are known to vary in the tissue/cell type and in a species-specific manner^35,42–44^. Given the possible roles and transcriptomic alterations in oligodendrocytes and astrocytes following stress and excess GC^45,46^, major differences between our and the previously reported GC-primed genes likely result from differences in the type of samples used namely *in vitro* human neurons versus zebrafish whole-brain samples containing neuronal and non-neuronal cells.

Interestingly, we identified a large set of upregulated DEGs were enriched in DNA metabolism, various RNA processing processes, and posttranslational modifications as well as epigenetic modifiers including DNA methyltransferases. For example, we have identified a subset of upregulated dGC-primed RNA splicing factors that are involved in U1, U2 or U12-dependent mRNA splicing, including *rbmx2, rbm5, srrm1, rbm25a/b, luc7l,* and *rnpc3*^47–50^. Also, we observed a downregulation of genes encoding dGC-primed RNA-binding proteins, such as *elavl4, qkib, celf2, celf5a, celf5b, nova1,* and *pum2*, which may play important roles in the nervous system^36–41^. Moreover, the most commonly identified transcript variances between bPAC+ vs bPAC- were alternative transcription start sites (ATSSs) and ATTSs. In the context of tissue-dependent exon usage, it was reported that the alternative transcription start and termination sites played an important role^51^. Therefore, the identified ATSSs and ATTSs associated with dGC-primed genes may be regional or cell-type dependent alterations, indicating that dGC-exposure may be able to prime genes in a cell-type or regional-specific manner. Furthermore, alternative donor sites are involved in exon definition^52^, repressor-activator competition^53^ as well as psychiatric disorders^54,55^. In a similar context, a recent study reported that the administration of dexamethasone, an activator of the GR, also results in alterations to the profile of alternatively spliced genes^56^. Since an increasing body of evidence underscores the role of alternative splicing in diseases including psychiatric disorders^54,57^, further studies like using long-read sequencing may reveal an even greater number of alternative splicing events induced by exposure to dGC. Collectively, dGC-exposure may increase the risk of dysregulation of gene expression in the later life stage by disturbing epigenetic regulations and RNA processing in adulthood.

Our results indicate that LD-exposure alters *nr3c2* (MR) and *crhr1* expression specifically in dGC-exposed animals while the expression of *nr3c1* (GR) is not altered. While GCs can bind both GR and MR in the brain, where many cells express both types of receptors, MR binding typically occurs at low levels of GC concentration, such as during resting state. In contrast, the lower affinity GR is usually occupied only under high GC concentration, such as under stress or at circadian peak^58^. Accordingly, MR and GR operate coordinately in a complementary fashion in response to environmental demands and an imbalance of MR/GR ratio is predicted to compromise the initiation and termination of the stress response leading to HPA axis dysregulation and impaired behavioral adaptation^59–61^. In this regard, it will be interesting to determine the specific site of MR:GR expression imbalance upon LD exposure in our model and assess their potential roles in affected behaviors in bPAC+ animals.

As early exposure to excess GC is considered a global risk factor for the development of mental disorders, our behavioral pipeline was tailored to assess performance in key behavioral manifestations rather than disease-specific tasks. bPAC+ fish did not show defects in aspects of perception as they can recognize visual and olfactory stimuli in the form of social cues and food pellets. Interestingly, bPAC+ fish exhibited a reduction in the maintenance of social interactions compared to the wild type, but not to bPAC-. We hypothesize that the attenuation of social interactions could potentially be influenced by paternal epigenetic memories resulting from developmental exposure to GCs. These effects should not be underestimated as they could serve as a significant contributory factor to the emergence of divergent social interaction patterns in adulthood. In previous zebrafish ELS studies, diminished shoaling behavior was observed following early-life social isolation and was associated with both motivational deficiencies linked to dopaminergic systems^62^ and increased sensitivity to stimuli which was reversed by anxiolytic treatment^63^. Impairments in social behavior or social withdrawal are common among different psychiatric disorders, including autism^64^ and schizophrenia^65^, and have been described in rodent models of ELS in which deficits in social interaction in the absence of social recognition and approach have been observed^66^. Further, bPAC+ fish showed deficits in fear learning compared to wild type using a Pavlovian conditioning paradigm that pairs electric shock with the color green. It is possible that perceived salience of electric shock as a stressor rather than fear learning is diminished in bPAC+. However, we consider this possibility less likely as bPAC+ can respond robustly to acute stress presented in the form of LD. This phenotype is consistent with what has been reported in rodent ELS models, which showed deficits in fear learning, learning impairment, and decreased cognitive flexibility^67–69^. Differences in how zebrafish learn fear association are dependent on whether they are reactive or proactive responders to stress, raising the question of how bPAC+ fish’s response fits into that spectrum^70^.

Recently, gene catalogues of neuropsychiatric disease-associated genes following stressful events including early life stress were reported based on gene expression network, eQTLs and GWAS^32,71,72^. The majority of dGC-primed genes which are associated with schizophrenia, bipolar disorder, depressive disorder, and autistic disorder-associated showed an exaggerated transcriptional response to stress in adult bPAC+ brains. Those DEGs were overrepresented in synaptic signaling processes including genes for GABA receptors (*gabrg2*, *gabbr1a*, *gabbr1b*, *gabra1*, *gabra2a*, *gabra4*, *gabrg2*), glutamate receptors and transporter (*grm1b*, *grm3*, *grm5b*, *grik3*, *grin2b*, *slc17a7a*), dopamine receptors (*drd2a*), cholinergic receptor (*chrm2a*), opioid receptor (*pnoca*), and neurexins (*nlgn1*, *nlgn4xb*). In a human study, genetic variants associated with GR-mediated immediate transcript response were able to predict risk for psychiatric disorders including depression^73^. Consistent with this, our findings suggest developmental GC exposure leads to differences in GR-induced transcriptional activation in adult brains and may mediate the risk for psychiatric disorders by altering a network of neuronal signaling-related stress-sensitive genes.

In conclusion, we identified highly exaggerated transcriptional response in animals with exposure to a high level of GC during development. However, further investigation is required to reveal the consequence of altered epigenetic modulators and a direct mechanistic link between the identified dGC-primed genes and alterations in regulatory mechanisms.

We propose that the brain DEG set identified here will be useful for guiding future studies dissecting the mechanisms underlying developmental GC-exposure and alteration of behaviors and neuropsychiatric disease susceptibility in adulthood. The zebrafish model presented here provides an opportunity to dissect the long-term effects of endogenous GC elevation during development and can be leveraged in future studies to identify the critical period, intensity, and duration of developmental GC exposure as well as epigenetic regulatory mechanisms including RNA processing and DNA methylation that leads to adult dysfunction.

## Methods and materials

The detailed protocol for the generation of transgenic animal, behavioral tests in this study is available as step-wise online methods^74^.

### Zebrafish husbandry and maintenance

The tübingen (TU) strain and transgenic line, *Tg(star:bPAC-2A-tdTomato)^uex300^* were kept at 28°C on a 12:12 hour light/dark cycle and housed at a maximum density of 5 fish/L. For experiments with adult fish, only females were used. All animal procedures were carried out in compliance with the ethical guidelines of the national animal welfare law and approved by relevant authorities (Landesuntersuchungsamt Rheinland-Pfalz, Germany, Project number 23 177-07/G20-1-033 or UK Home Office PPL number PEF291C4D).

### Sample collection

All samples were collected within a 2-hour window in the morning (08:30 to 10:30) without feeding.

### Whole-body Cortisol assay

The competitive cortisol assay (Cisbio HTRF® Cortisol Kit, 62CRTPEG) was performed following the manufacturer’s protocol. ELISA signal was detected by CLARIO star plate reader (BMG Labtech, Ortenberg, Germany).

### Behavior tests

Social behavior tests and acute stressor delivery using looming dot (LD) presentation were carried out as described in Cook *et al.*^21^. Basal locomotion, food consumption, and fear conditioning assay were performed following the protocol in online methods^74^. For all behavioral tests, female adult zebrafish (6-9 months old) were moved into a designated behavior experimental room one week prior to testing and housed in groups of 10 fish in 3L tanks.

### RNA preparation and quantification

Each sample for the larval stage (6 and 13 dpf) contained 25-30 larval whole brains whilst for the adult stage (120 dpf) three brains constituted one sample. We dissected the larval whole brains (at 6 or 13 dpf) in the RNAlater® (AM7021, Ambion, USA) after overnight incubation and the adult brains (at 120 dpf) were dissected in PBS on ice and then snap freeze in liquid nitrogen. Samples were kept at −80°C until further processing. Larval and adult samples were completely homogenized in RNA Lysis buffer from the Quick-RNA miniprep Kit (R1055, Zymo Research, Irvine, CA, USA) using a pestle and micro-tube homogenizer (for 30 sec) or TissueLyser LT (QIAGEN, Dusseldorf, Germany, at 25 Hz for 1 minute and then 15 Hz for 2 minutes), respectively. Total RNAs were isolated by following the manufacturer’s protocol and keep it at −80°C until further use. Transcripts were quantified using real-time qPCR and next-generation mRNA sequencing. cDNA was synthesized using a High-Capacity RNA-to- cDNA™ Kit (4387406, Applied Biosystems, USA) following the manufacturer’s protocol. The real-time qPCR was performed with PowerUp™ SYBR™ Green Master Mix (A25778, Applied Biosystems, USA) and specific primers. Primer information for qPCR is described in online methods^74^. mRNA-seq library preparation and sequencing were performed by TRON gGmbH (Mainz, Germany) using the Illumina NovaSeq6000. Briefly, construct paired-end TruSeq Stranded mRNA libraries (Illumina, CA, USA) and sequence it for over 20M of 50 bp reads/sample. A total of 60 samples was sequenced, consisting of 5 biological replicates at four different time points (6, 13, 120 dpf, and acute-stressed at 120 dpf) for each genotype (wild type, bPAC+ and bPAC-).

### DNA preparation and quantification

Whole brains were collected from 9-month-old bPAC+ and wild type fish. 6 individual brains per group were used for DNA isolation using DNeasy® Blood & Tissue kit (#69504, QIAGENE) following the manufacturer’s protocol. Purified DNA was sequenced using the Oxford nanopore sequencing system. Ligation sequencing libraries were prepared from 1000 ng DNA using the Ligation Sequencing kit SQK-LSK114 (ONT, Oxford Nanopore Technologies Ltd., Oxford, UK) following the manufacturer’s protocol. The clean-up step after adapter ligation was intended to size-select fragments and was done with Long Fragment Buffer. The library was loaded on a single R10.4.1 (FLO-PRO114M) flow cell and sequenced on a PromethION 24 device within 72 h. MinKNOW (v22.12.05) was used to supervise the initial sequencing run.

### Bioinformatic analyses

#### mRNA sequencing

RNA sequencing analyses were performed following the online methods^74^. Briefly, processed qualified sequencing reads by FASTQC^75^, fastp^76^ were mapped to the zebrafish reference genome assembly (GRCz11) with Ensembl annotation version 107 using HISAT^77^. We estimated the expression of transcripts using Stringtie^78^ and DEGs using edgeR^79^. Downstream and statistical analyses were performed with in-house R scripts on and Rstudio (Build 485)^80^. DEGs were defined with criteria; |FC| > 1.5 and FDR < 0.05 for the comparisons between bPAC+ vs. bPAC-, or |FC| > 2 and FDR < 0.01 for the comparisons between bPAC+ vs. wild type. Functional analyses were performed with the R packages including clusterprofiler^81^, gProfiler2^82^, enrichGO^81^ and DisGeNET^32^. Protein-protein interaction networks were constructed and visualized by using the String database^83^ and Cytoscap^84^. Alternative splicing and isoform switches were analyzed by using IsoformSwitchAnalyzeR^85^.

#### Long Read Sequencing

The sequenced reads quality assessment was done using pycoQC^86^. The reads from each sample were base-called using Guppy (v6.4.2, Oxford Nanopore Technologies Ltd) and mapped to the *Danio rerio* (GRCz11) genome using minimap2 aligner^87^. 5mC CpG modified bases were determined using the modbam2bed tool (Oxford Nanopore Technologies Ltd).

#### Differential methylation analysis

Differential methylation analysis between control and stress samples was done using DSS Bioconductor package^88^ with p < 0.05. Bases were considered as differentially methylated if FDR < 0.05 and absolute DNA methylation difference between two groups larger than 25%.

Genomic coordinates of exon, intron, promoter, and intergenic region for *Danio rerio* genome were obtained from the UCSC genome browser. Promoters were defined as upstream 800bps and downstream 200bps from TSS. Differentially methylated cytosines were annotated based on genomic positions into exons, introns, promoters, and intergenic regions using the Genomic Ranges findOverlaps function^89^. Methylation browser tracks were created using deeptools bamCoverage utility^90^. The variant calling was performed using Clair3 package^91^.

### Code availability

The detailed data processing and R scripts for bioinformatic analysis are available at the GitHub repository (https://github.com/minkechoi/tx_star-bPAC_brain).

### Data availability

All sequenced reads for RNA-seq were deposited in European Nucleotide Archive (ENA, PRJEB53713).

### Statistics

Statistical analyses were performed using R and Prism 9 (Graphpad Software Inc, San Diego, CA, USA). A detailed description of the statistical analyses is provided in online methods^74^

## Acknowledgment

This project was funded by the German Federal Office for Education and Research (BMBF) grant number 01GQ1404 and Mireille and Dennis Gillings Foundation Grant to SR, Basic Science Research Program (2020R1A6A3A03037828) of National Research Foundation (NRF) of Korea to MC and Institutional Strategic Support Fund 3 scheme (ISSF3) to Translational Research Exchange @ Exeter (ISSF3-TREE-Choi, 2022) funded by Wellcome Trust to MC. We acknowledge the support of TRON gGmbH (Mainz, Germany) for the RNA-sequencing, Aquatic Resources Center (University of Exeter, UK), and Kathrin Domdera (University of Mainz, DE) for expert fish care, and High-Performance Computing (HPC) facility (University of Exeter, UK) for the server for informatic analyses.

## Author contributions

SR conceived the project. MC and SR designed the project. MC acquired and analyzed omics data. KM and ML generated and analyzed ONT-seq data. AC acquired and analyzed behavior data. HE, and AT contributed to reagent generation or data acquisition. MC, SR, and AC drafted the article. MC, HE, and SR critically revised the manuscript. All authors approved the submission.

