## Supplemental Figures for "Lifelong molecular consequences of high Glucocorticoids exposure during development"

Supplementary Figure 1. The Strategy of Animal setup for experiments. S
Supplementary Figure 2. DEG identification and clustering.
Supplementary Figure 3. nr3c2 and crhr1 expressions were differentially regulated in bPAC+ following LD.
Supplementary Figure 4. Comparison between bPAC+ post-LD genes and DEGs identified in the human hippocampal progenitor cell line.
Supplementary Figure 5. Expression trajectories of DNA methylation modulator genes.
Supplementary Figure 6. Identification of differentially methylated CpGs (DMCs).

Supplementary data File 1. Volcano plots for all pairwise comparisons.
Supplementary data File 2. Switch plots for alternative splicing variants.

Supplementary Table 1. List of DEGs.
Supplementary Table 2. Results of GO enrichment test for DEGs.
Supplementary Table 3. List of Adult LD-DEGs.
Supplementary Table 4. Results of GO enrichment test for GC-primed DEGs.
Supplementary Table 5. List of disease-associated GC-primed DEGs.
Supplementary Table 6. List of overlapping genes with human GC-primed genes.
Supplementary Table 7. List of DMCs on mutual GC-primed genes associated regions.
Supplementary Table 8. List of epigenetic modifiers in LD-DEGs.
Supplementary Table 9. List of significant alternative splicing variants.

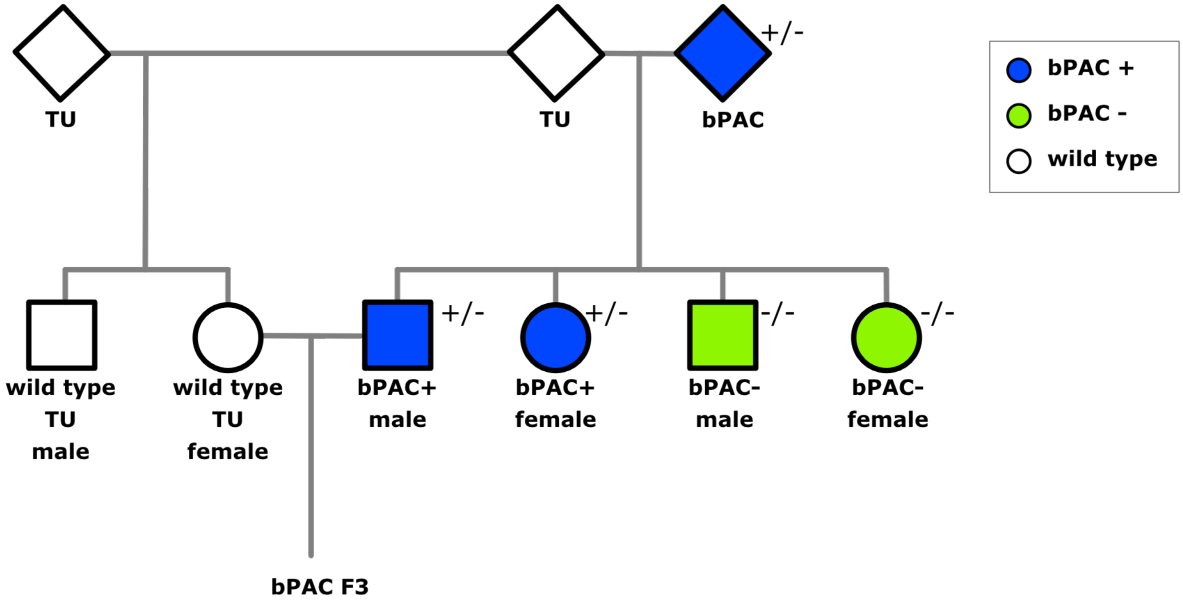


**Supplementary Figure 1. The Strategy of Animal setup for experiments.** bPAC+ transgenic fish line was maintained by outcrossing bPAC+ with wild type (TU). bPAC+ and bPAC- larvae were screened based on the tdTomato signal in interrenal gland at 4 days post-fertilization (dpf). However, for cortisol measurements during early developmental stages prior to onset of the tdTomato expression, we utilized progenies of homozygous bPAC+ crossed to wildtypes. Adult brain samples were collected from females.

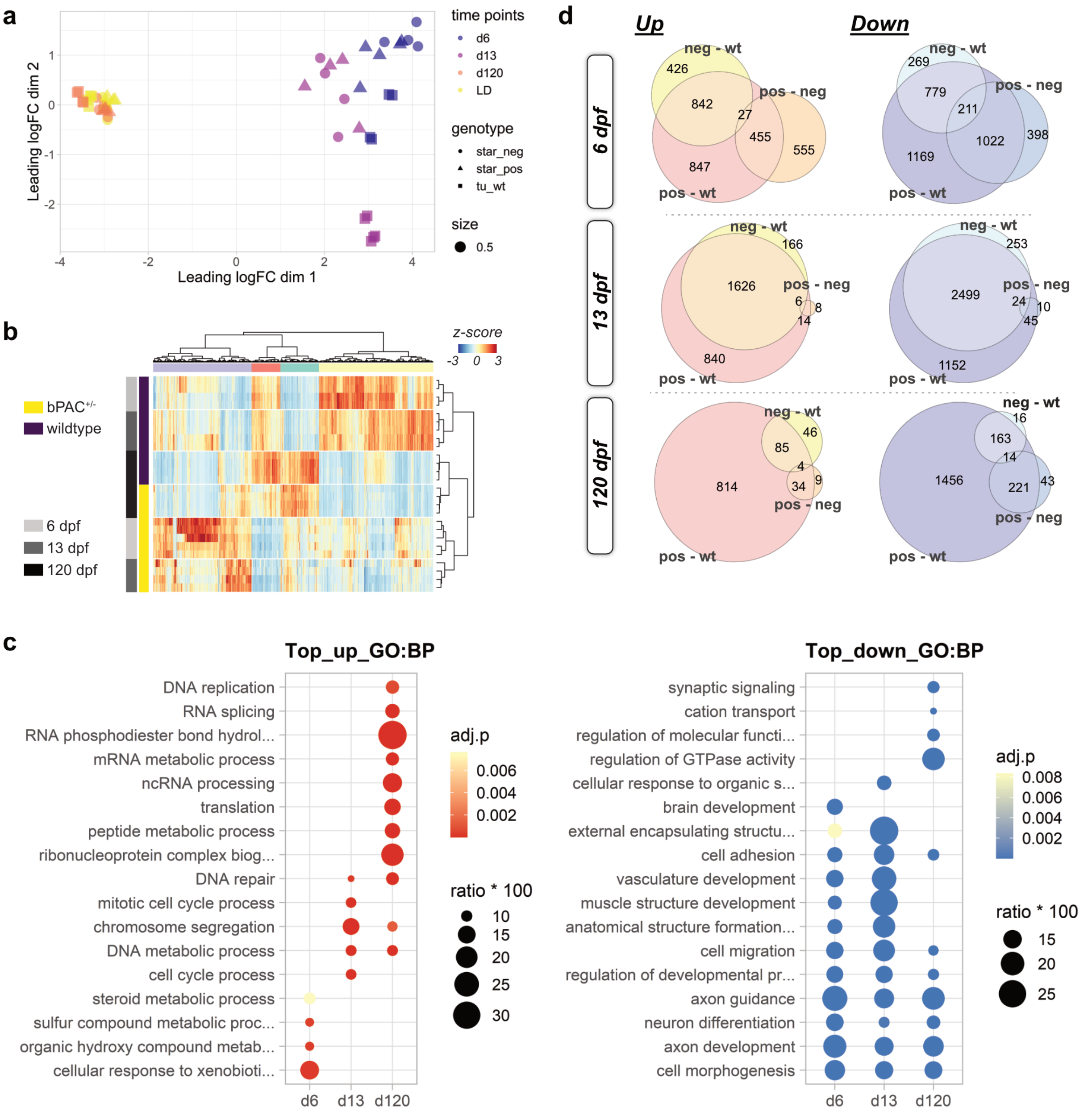


**Supplementary Figure 2. DEG identification and clustering.** (a) MDS plot for RNA-seq data. Global profiles of samples were clustered by time and genotype. Profiles of bPAC+ and bPAC- were more similar to each other than to those of wild type fish. (b) Read counts of genes by genotype and time points are represented by z-score in the heat map using bPAC+ and wildtype samples. Bars on the left side indicate the genotype of fish and time points. (c) Top GO terms for biological processes enriched in upregulated (left) or downregulated (right) DEGs for the comparisons between bPAC+ and wild type. d6: 6 dpf, d13: 13 dpf, d120: 120 dpf, wt: wild type, FC: fold change, GC: glucocorticoid, adj.p: adjusted *P*-value, ratio: identified DEGs/the number of genes included in the term. Complete DEG-enriched GO terms are described in S. Table 2. (d) Venn Diagram for up- and down-regulated DEGs among wild type, bPAC- and bPAC+ by time.

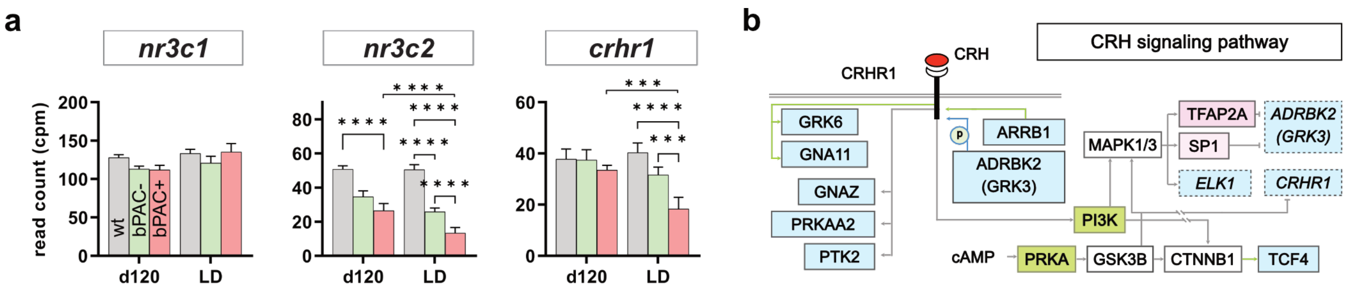


**Supplementary Figure 3. *nr3c2* and *crhr1* expressions were differentially regulated in bPAC+ following LD.** (a) Expression levels of *nr3c1*, *nr3c2*, and *crhr1* before and after LD exposure. The asteria is represent those comparisons that are statistically significant with |FC|>1.5. **=P<0.05*, ***=P<0.01*, ****=P<0.001*, *****=P<0.0001*. Tukey's multiple comparisons test followed Restricted maximum likelihood analysis. Error bars indicate mean+SD. (b) Following the LD exposure, several homologs in downstream of CRHR1 were differentially expressed in bPAC+ brains. Boxes colored light blue and pink indicate downregulation and upregulation of its transcripts, respectively. The green-colored boxes represent enzyme complexes. The green and grey lines showed known and unknown interactions, respectively. Solid and dotted outline of boxes represent protein and transcript, respectively. Pathway was sourced by WikiPathways (https://www.wikipathways.org/instance/WP2355) (Martens, et al., 2021).

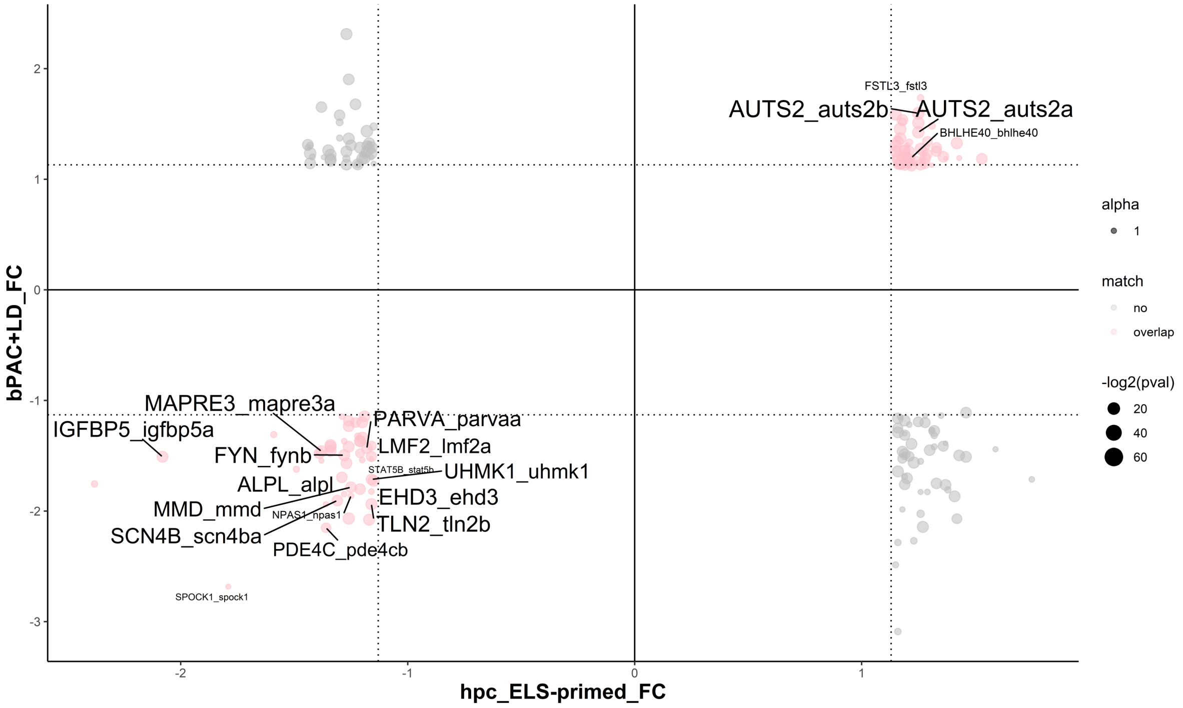


**Supplementary Figure 4. Comparison between bPAC+ post-LD genes and DEGs identified in the human hippocampal progenitor cell line.** DEGs mapped to long-lasting differentially methylated cytosines from a GC-exposed human hippocampal progenitor cell line study (Provençal et al., 2020) were compared to bPAC+ post-LD DEGs (i.e. bPAC+/bPAC- post-LD comparison). X-axis represents FC of post-LD DEGs in bPAC+, y-axis represents FC of DEGs in GC-exposed mature neurons from a GC-exposed human hippocampal progenitor cell line. Labeled genes indicate GC-primed genes in both studies. Size of text for gene names indicates significance, -$\text{Log}_{10}$(sqrt(P-values from studies)). Gene names are labelled as human gene_zebrafish gene.

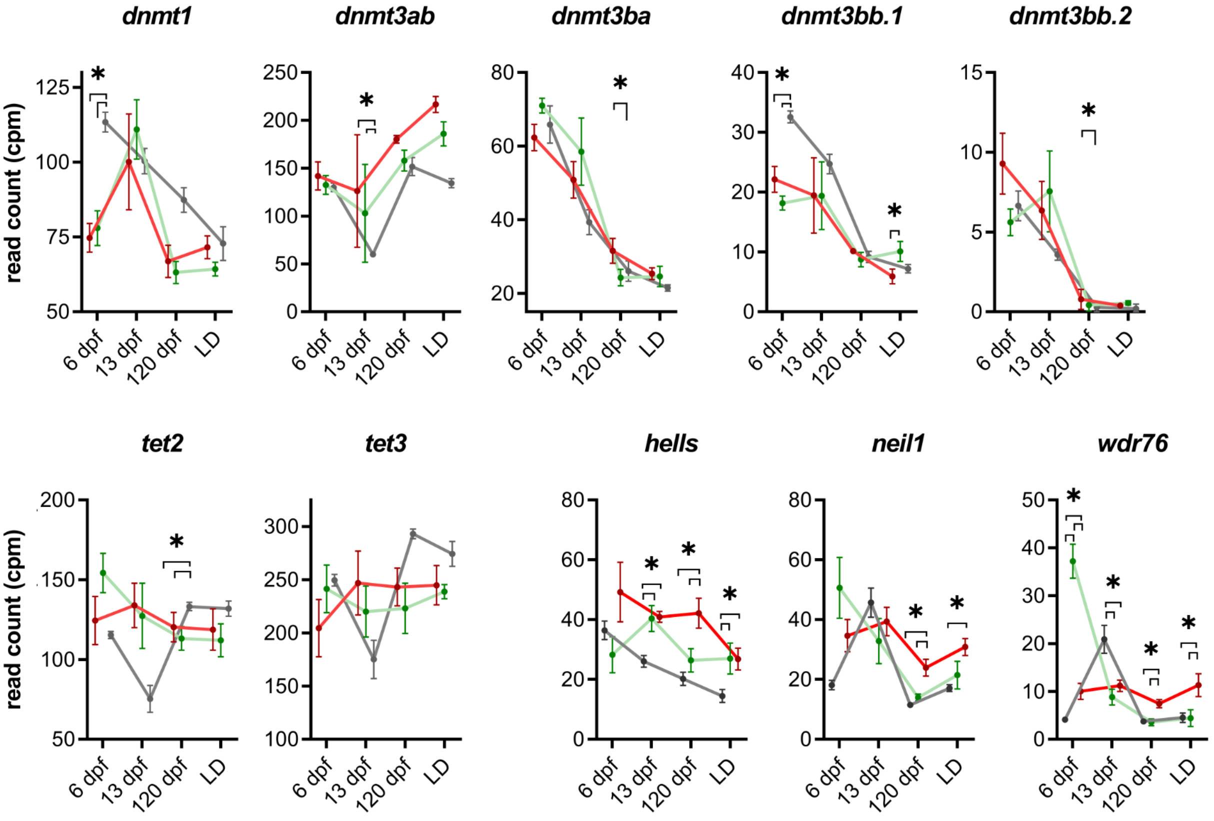


**Supplementary Figure 5. Expression trajectories of DNA methylation modulator genes**. 5mC writers (*dnmts*), 5mC erasers (*tets*), 5hmC readers (*hells, neil1, wdr76*). red line: bPAC+, green line: bPAC-, grey line: wild type. The asteria mark indicates significant differential expression (FDR < 0.05, |FC| < 1.5).

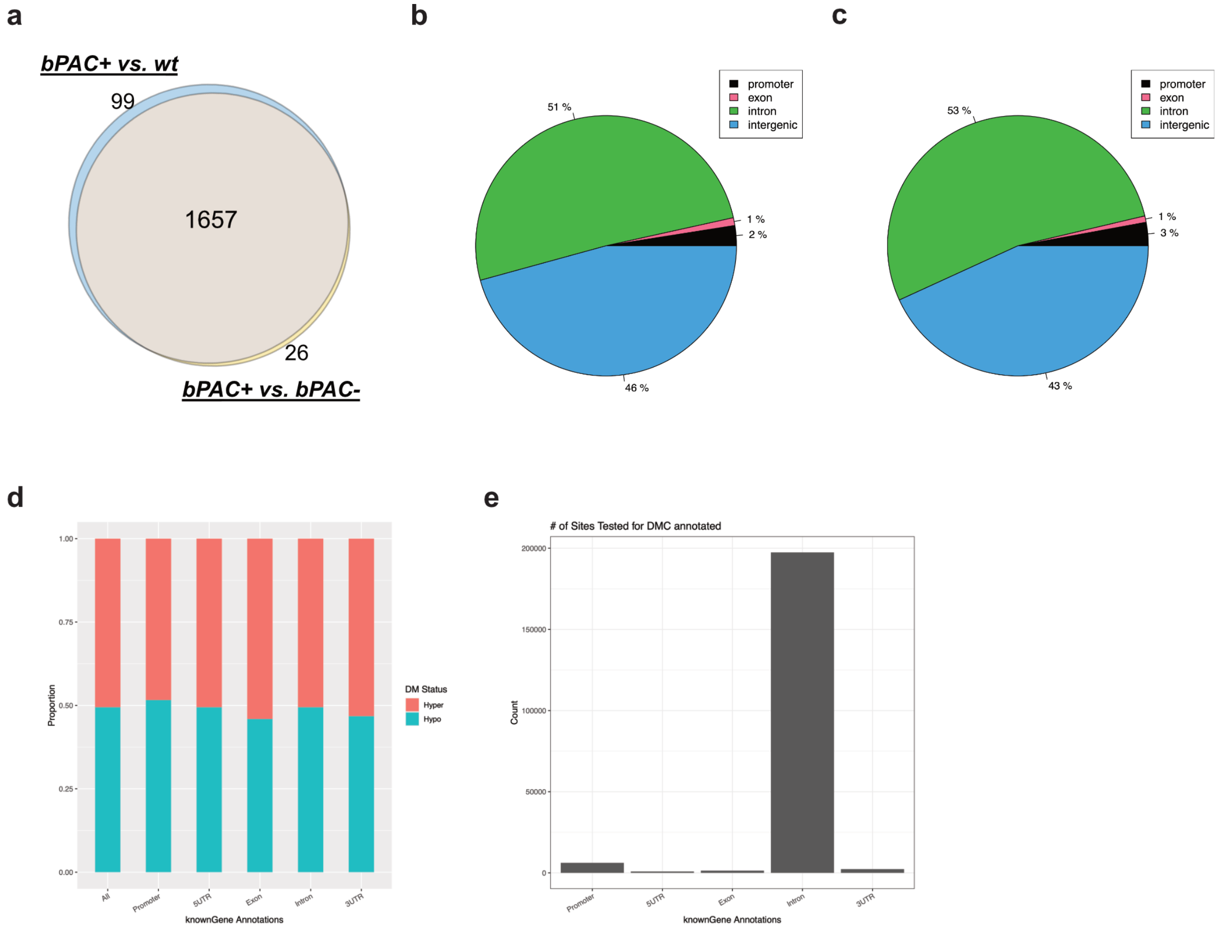


**Supplementary Figure 6. Identification of differentially methylated CpGs (DMCs).** (a) Venn diagram showing dGC-primed genes from two pairwise comparisons, bPAC+ vs. wild type and bPAC+ vs. bPAC-. Distribution of Hypo- (b) and Hyper-methylated (c) DMCs according to the gene regions represented by Pie chart. (d)Proportion of Hypo and Hyper-methylated DMCs according to the gene regions represented by and Bar charts. (e) The number of identified DMCs according to the gene regions.
